# The SOD-2 protein is the single active SOD enzyme in *C. elegans*

**DOI:** 10.1101/2020.08.12.247973

**Authors:** Lourds M. Fernando, Silwat Adeel, Mohammed Abul Basar, Anna K. Allen, Atanu Duttaroy

## Abstract

The nematode *C. elegans* has a contingent of five *sod* genes, one of the largest among aerobic organism. Earlier studies revealed each of the five *sod* genes is capable of making perfectly active SOD proteins in heterologous expressions systems therefore none appears to be a pseudogene. Yet deletion of the entire contingent of *sod* genes fails to impose any effect on the survival of *C. elegans* except these animals appear more sensitive to extraneously applied oxidative stress condition. We asked how many of the five *sod* genes are actually active in *C. elegans* through an in-gel SOD activity analysis. Here we provide evidence that out of the five genes only the mitochondrial SOD gene is active in *C. elegans*, albeit at a much lesser amount compared to *D. melanogaster* and *E. coli*. Mutant analysis further confirmed that among the mitochondrial forms, SOD-2 is the only naturally active SOD in *C. elegans*.

## 1. Introduction

Aerobic organisms utilize Superoxide Dismutase (SOD) enzymes to scavenge and catabolize partially reduced Oxygen molecules (superoxide radicals; O_2_^−^) into Hydrogen Peroxide (Fridovitch, 1995). Superoxides are copiously generated in the electron transport chain from aerobic respiration. In different aerobes, SOD enzymes entrap superoxide radicals with the help of a variety of redox active transition metals such as Cu, Mn, Fe or Ni. Despite all SODs performing one single task, that is to scavenge superoxide radicals, specific SOD enzyme activities are noticeably limited to either the mitochondrial matrix, cytoplasm, inter-membrane space, or to the extracellular environment (Weisiger and Fridovich 1973; Missirilis et al., 2003; Muller et al., 2004). Loss-of-function studies in different species have established the variable requirement of the SOD protection system because the spectrum of defects appearing in *SOD* mutants are wide ranging. In general, lack of mitochondrial SOD in vertebrates cause neonatal lethality accompanied by signs of mitochondrial damage, cardiomyopathy and neuropathology (Lebovitch et al., 1996; Li et al., 1995). Early lethality is also evident in Drosophila mitochondrial SOD (SOD2) loss of function condition (Kirby et al., 2002; Duttaroy et al., 2003) with evidence of neuropathology (Paul et, al., 2007), loss of mitochondrial aconitase activity and reduced ATP production in the muscle (Godenschwege et al., 2009).

On the other hand, loss of Cu-ZnSOD enzyme (aka SOD1) causes more subtle effects particularly in higher animals. For example, mice lacking Cu-Zn SOD activity exhibit no effect on longevity although a multitude of other damages are reported (Valentine et al., 2005; Culotta et al., 2006). Drosophila *Cu-ZnSod* null mutants survive albeit with a shorter adult life span (Phillips et al., 1989). Finally, bacterial and yeast mutants of *Sod1* and *Sod2* are hypersensitive to hyperbaric Oxygen concentration, and exhibit numerous biochemical defects in these mutants (Culotta, 2000).

The nematode *C. elegans* is a free-living soil dwelling aerobe traditionally used as a workhorse in studies related to aging, cell death, and developmental biology (Corsi et al., 2015). Homology analysis established five *Sod* genes in *C. elegans*, possibly one of the largest contingents of SOD proteins among aerobic organism (Landis and Tower, 2005; Worm Base). *sod-1, sod-2* and *sod-4* in *C. elegans* represent the traditional cytoplasmic, mitochondrial and extracellular forms of SODs respectively, while *sod-3* and *sod-5* are recognized as additional and minor mitochondrial and cytoplasmic SODs (Doonan et al., 2008). *sod-1* and *sod-2* mRNAs represent the bulk of the *sod* RNAs (∼76% and 18% respectively) within the total SOD mRNA populations whereas the rest of the three *sod* mRNAs are represented between 5-0.5% of the total (Doonan et al., 2008). So, in *C. elegans* all *sod* genes are expressed but differentially.

Are these extra copies of *sod* purposefully acquired to gain additional protection against some pro-oxidative environment or are most of these *sod* genes evolutionary relics? Mutational analyses have already provided a striking answer as *C. elegans* carrying quintuple deletions of all *sod* genes survive as long as the wild type control worms (Van Raamsdonk and Hekimi, 2012). Additionally, no tissue damage or loss of mitochondrial proteins was evident due to the lack of a complete SOD protection system. The only noticeable change seems to be these animals are hypersensitive to different kinds of oxidants (Van Raamsdonk et al., 2009). However, *C. elegans sod* genes are not evolutionary relics because the *sod-1, sod-2, sod-3* and *sod-5* genes can make perfectly active SOD proteins in heterologous systems like bacteria and yeast (Hunter et al., 1997; Jensen and Cullota, 2005), therefore no coding sequence deficiencies are expected in these genes like in a pseudogene. Also, SOD enzyme activity assays support that *C. elegans* do carry active SOD enzyme(s) (Doonan et al., 2008; Ramsdonk and Hekimi, 2012) but which one or how many of those five SOD proteins are naturally active remains an open question.

## 2. Materials and Methods

### C. elegans strains and culture conditions

The following *C. elegans* strains were used in this work: Bristol strain N2, CB1370 [*daf-2(e1370) III*], MQ1776 [*sod-2(ok1030) I; sod-5(tm1146) sod-1(tm783) II; sod-4(gk101) III; sod-3(tm760) X*], VC433 [*sod-3(gk235) X*], and FX776 [*sod-1(tm766) II*]. All strains were grown and maintained under standard conditions at 20**°**C.

### C. elegans protein extraction

Native *C. elegans* protein lysate was prepared by washing the appropriately staged worms off MYOB plates grown at 20**°**C with 1XPBS into a 1.5ml microcentrifuge tube, pelleting the worms with gentle centrifugation, and homogenizing the worm pellet on ice for 10mins. The tubes were centrifuged at 16000Xg at 4**°**C for 15mins and the supernatant was aspirated carefully avoiding the floating lipid layer. Bradford assay was performed to measure the protein concentration.

### In-gel SOD activity assay

For in-gel SOD activity assay total protein extracts (60μg/lane) were loaded into a 12% non-denaturing gel. Once gel running was completed they were soaked in 1.23 mM NBT (Nitro-blue Tetrazolium), rinsed briefly, and soaked in 100 mM potassium phosphate buffer (pH 6.8) containing 28 mM TEMED and 2.8 mM riboflavin used as nascent free radical donors in a photochemical reaction that converts NBT to blue formazan. Gels were exposed to white light for development of signals.

## 3. Results and Discussion

To answer how many and which particular SOD protein is active we performed *in situ* SOD gel analysis using preparations of native protein extracts from *C. elegans* and compared this with *Drosophila* and *E. coli* extracts. Following electrophoresis, gels are incubated in NADPH substrate solution which serves as the electron donor. Presence of active SOD enzyme(s) in its native state clears the blue formazon locally and appears as bright, white bands in the gel (Fig. 1). Starting with *Drosophila*, two active SOD enzymes are identified (Fig. 1A) with the top band being the mitochondrial SOD (MnSOD or SOD2) and the lower band representing the cytoplasmic SOD (Cu-ZnSOD or SOD1) (Phillips et al., 1989; Duttaroy et al., 2003). As expected, the *E. coli* extract shows three forms of SOD enzymes MnSOD, Fe-SOD, and the Cu-ZnSOD (Figure 1A). On the other hand, wild type *C. elegans* (N2) extract shows only a single SOD enzyme band indicating that it carries no more than one active form of a SOD enzyme (Figure 1A). Two things are notable with respect to the active *C. elegans* SOD enzyme: (1) it lines up perfectly with the MnSOD/SOD2 isoforms of *Drosophila* and *E. coli*, and (2) *C. elegans* SOD activity appears as a much less intense band in comparison to the other organisms despite loading equal amounts of protein extracts (60 μg/lane) per lane. This leads us to hypothesize that either *C. elegans* makes much less SOD enzyme or that the enzyme itself is less active in this organism compared to the *Drosophila* and *E. coli* SODs. Attempts to enhance the SOD band intensity through longer incubation of the gels in NADPH substrate or by using greater amounts of substrate failed to enhance the intensity of this SOD band (not shown). Further research should establish if *C. elegans* SOD is either expressed at a lesser amount or if is it less active relative to other organisms.

**FIGURE 1:**
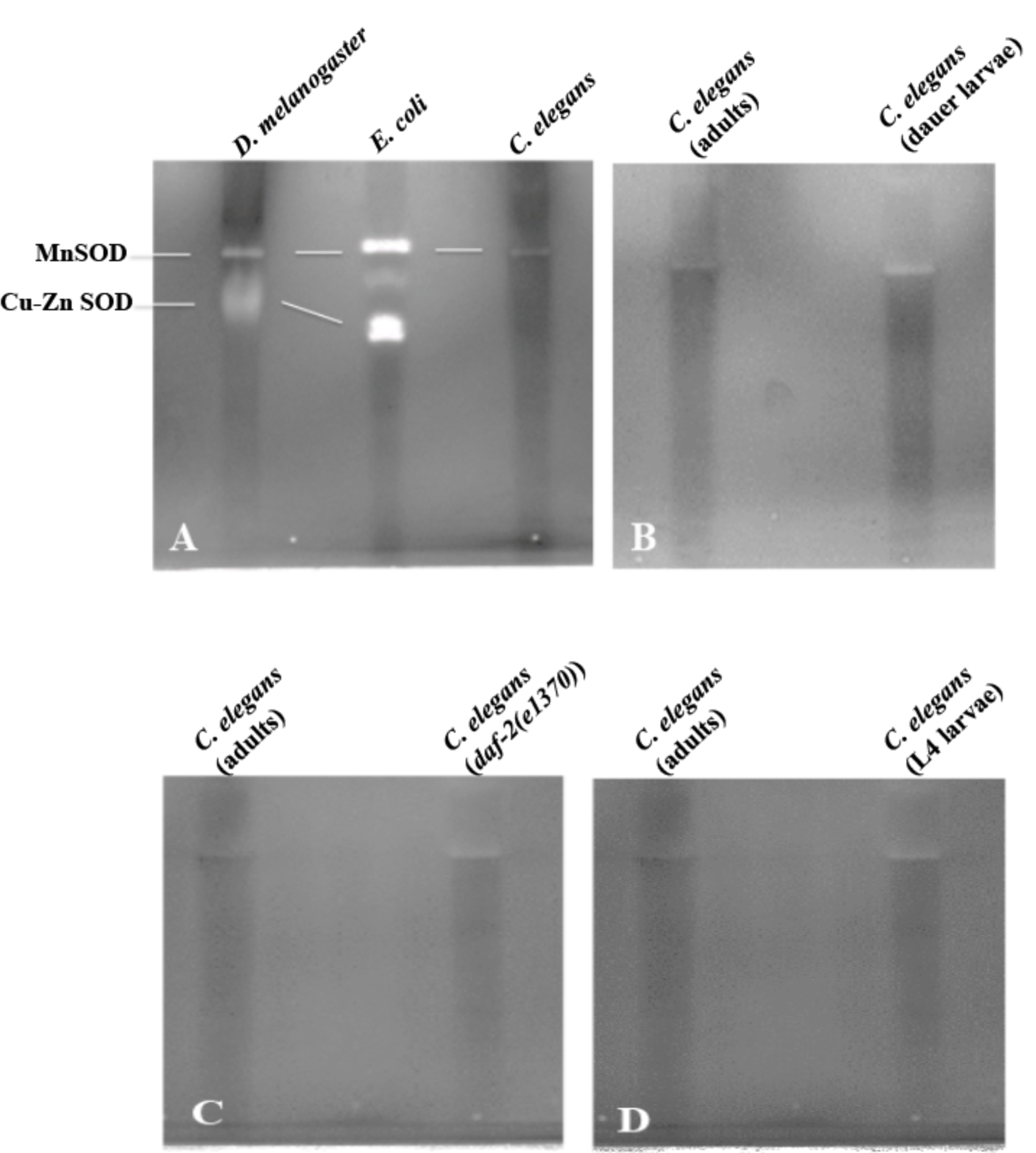
*In situ* SOD enzyme profiles in three different species. (A) Comparison of SOD activity profiles in *D. melanogaster, E. coli* and *C. elegans. C. elegans* has no more than one active SOD enzyme. (B, C, and D) The dauer larval stage, *daf-2* mutant and L4 larvae all carry no more than one active form of SOD.

Since none of the five s*od* genes in *C. elegans* are pseudogenes as each is capable of making perfectly active SOD protein in a heterologous system (Hunter et al., 1997; Jensen and Cullota, 2005), then why is there only one naturally active SOD enzyme? We tested a few possibilities such as some of the SOD enzymes might be made and active early during the growth and development of the worm (Jenese and Cullota, 2005), or some SODs are active only under high stress condition. However, no additional SOD enzymes appear active in the L4 larval stage protein extracts compared to adult extracts (Fig 1C) nullifying the possibility of a developmental activation of certain SOD enzymes (Figure 1C). Similarly, *C. elegans* dauer arrested larvae also show no more than one active SOD enzyme (Fig 1B). It has previously been reported that a *daf-2* mutant exhibits overproduction of *sod* mRNA (Honda and Honda, 1999). In our assays we observe a slightly more intense SOD band in the *daf-2* mutants supporting the reported increase in *sod* mRNA. Therefore, taken together it seems that only one SOD enzyme is naturally active throughout the lifespan in *C. elegans*.

The most obvious question appears to be which one of the five SOD proteins is the solely active enzyme in *C. elegans?* We pointed out earlier that the single active *C. elegans* SOD enzyme lines up nicely with the mitochondrial SOD2 band of *Drosophila* and *E. coli* (Figure 1A). This means either one of the two mitochondrial SOD proteins (SOD-2 or SOD-3) or maybe both are represented in this single SOD active enzyme band. Apparently these two SOD proteins arise from a gene duplication event (Hunter et al., 1997) and the predicted molecular weights of *C. elegans* SOD-2 and SOD-3 are 24.5 and 24.6kD respectively (Table 1) which matches very closely with the *Drosophila* and *E. coli* SOD2 proteins that are 24.7 and 24.5kD respectively (Table 1). The only other SOD that can match the sizes of SOD-2 and SOD-3 is SOD-4b isoform that weighs 23.3 kD (Table 1). But SOD4b with its large trans-membrane domain remains associated with the cell membrane (Fujii et al., 1998) and therefore it was eliminated with the membrane fraction during protein extraction. On the other hand, the cytoplasmic SODs like SOD-1, SOD-4 and SOD-5 in *C. elegans* are significantly different in size from the mitochondrial SODs weighing 18.7, 18.2 and 18.5kD respectively (Table 1). Yet the single SOD band in *C. elegans* was previously designated as the SOD-1 enzyme (Jenese and Cullota, 2005) so we used some *sod* mutants to resolve this issue. The quintuple *sod* knockout mutant worms [*sod-2(ok1030) I; sod-5(tm1146) sod-1(tm783) II; sod-4(gk101) III; sod-3(tm760) X*] (Van Raamsdonk and Hekimi, 2012) is lacking any active SOD enzyme (Fig. 2B Lane 2). However, the homozygous *sod-1* null mutant allele [*sod-1*(*tm776)*] and homozygous *sod-3* null mutant allele [*sod-3*(*gk 235)*] both are expressing the single SOD band like the wild type (Fig 2B). This leads us to conclude that the active SOD enzyme in *C. elegans* is the mitochondrial SOD-2 protein. In support of our conclusion, work by Doonan et al., (2008), previously showed wild type *C. elegans* expresses an abundant amount of mitochondrial SOD proteins including both SOD-2 and SOD-3, but very little amount of SOD-1 protein, although no true quantitative information was available (Doonan et al., 2008, Supplemental data).

**FIGURE 2:**
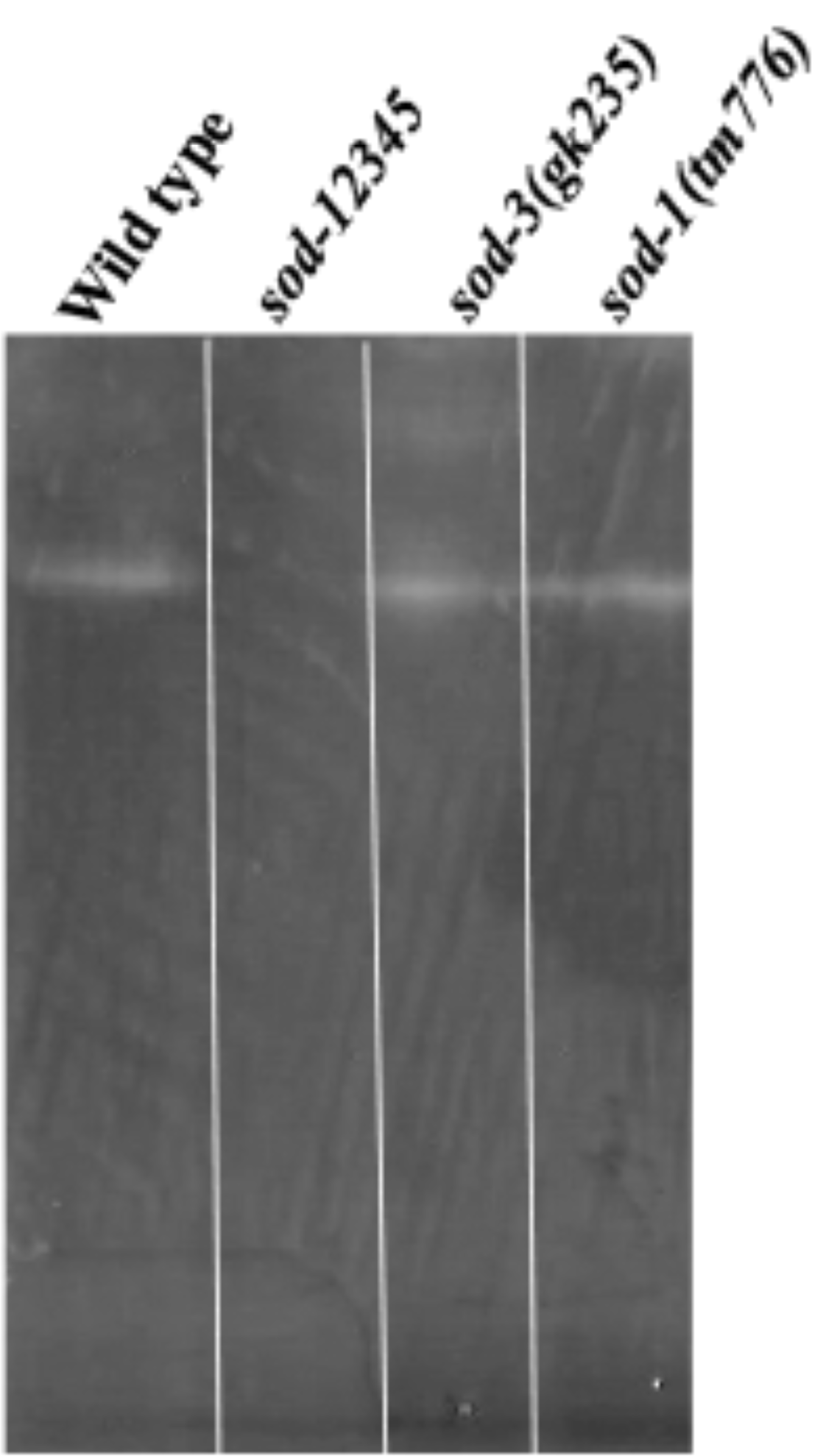
Active SOD activity profiles in different *C. elegans sod* mutants. (A) *sod-1(tm776)* is a *sod-1* null mutant which expresses the same active SOD as in wild type. (B) No active SOD appeared in the quintuple *sod* deletion line [*sod-2(ok1030) I; sod-5(tm1146) sod-1(tm783) II; sod-4(gk101) III; sod-3(tm760) X*], whereas (C) *sod-3(gk235)* which is a *sod-3* deletion mutant carries the single active form of SOD.

Our attempt to profile the active SOD enzymes in *C. elegans* led to the finding that the mitochondrial SOD-2 enzyme is the solely active SOD enzyme throughout the life span of this organism. Since the mitochondria is the main source of superoxide radicals in the cell due to the increased Oxygen metabolism in this organelle, keeping the mitochondrial enzyme in active state makes more sense for any aerobic organism. Although, why this lone SOD-2 enzyme appears relatively less active compared to other organisms remains unclear (Fig. 1A). Although active, the nonessential nature of this lone active SOD-2 enzyme is demonstrated as the *sod-2* deletion mutant positively influences the life span by surviving longer than the wild type and also exhibits other mutant phenotypes such as slow development, a low brood size, and slow defecation (Raamsdonk and Hekimi, 2009). Unlike any other aerobes, total abrogation of all SOD enzymes is tolerated by *C. elegans*, as well as it is completely resistant to hypoxic environment (Mehta et al., 2009). In fact, hypoxia imposes a positive influence on *C. elegans* life span. In sum, these observations challenge the necessity of SOD system as the primary defense mechanism towards ROS detoxification in *C. elegans*. The important question that remains is what has taken the place of the SOD system requirement in *C. elegans*?

## Acknowledgement

We thankfully acknowledge the National Science Foundation grant 1832026 and NIH grant R25 AG047843 awarded to AD. LMF and AKA are supported by a Department of Defense grant W911NF1810465 awarded to AKA.

